# A Probabilistic Programming Approach to Protein Structure Superposition

**DOI:** 10.1101/575431

**Authors:** Lys Sanz Moreta, Ahmad Salim Al-Sibahi, Douglas Theobald, William Bullock, Basile Nicolas Rommes, Andreas Manoukian, Thomas Hamelryck

## Abstract

Optimal superposition of protein structures is crucial for understanding their structure, function, dynamics and evolution. We investigate the use of probabilistic programming to superimpose protein structures guided by a Bayesian model. Our model THESEUS-PP is based on the THESEUS model, a probabilistic model of protein superposition based on rotation, translation and perturbation of an underlying, latent mean structure. The model was implemented in the deep probabilistic programming language Pyro. Unlike conventional methods that minimize the sum of the squared distances, THESEUS takes into account correlated atom positions and heteroscedasticity (i.e., atom positions can feature different variances). THESEUS performs maximum likelihood estimation using iterative expectation-maximization. In contrast, THESEUS-PP allows automated maximum a-posteriori (MAP) estimation using suitable priors over rotation, translation, variances and latent mean structure. The results indicate that probabilistic programming is a powerful new paradigm for the formulation of Bayesian probabilistic models concerning biomolecular structure. Specifically, we envision the use of the THESEUS-PP model as a suitable error model or likelihood in Bayesian protein structure prediction using deep probabilistic programming.

## I. INTRODUCTION

In order to compare biomolecular structures, it is necessary to superimpose them onto each other in an optimal way. The standard method minimizes the sum of the squared distances (root mean square deviation, RMSD) between the matching atom pairs. This can be easily accomplished by shifting the centre of mass of the two proteins to the origin and obtaining the optimal rotation using singular value decomposition [1] or quaternion algebra [2], HYPERLINK \l “bookmark2” [3]. These methods however typically assume that all atoms have equal variance (homoscedasticity) and are uncorrelated. This is problematic in the case of proteins with flexible loops or flexible terminal regions, where the atoms can posit high variance. Here we present a Bayesian model that is based on the previously reported THESEUS model [4]–[6]. THESEUS is a probabilistic model of protein superposition that allows for regions with low and high variance (heteroscedasticity), corresponding respectively to conserved and variable regions [4], [5]. THESEUS assumes that the structures which are to be superimposed are translated, rotated and perturbed observations of an underlying latent, mean structure **M**.

In contrast to the THESEUS model which features maximum likelihood parameter estimation using iterative expectation maximization, we formulate a Bayesian model (THESEUS-PP) and perform maximum a-posteriori (MAP) parameter estimation. We provide suitable prior distributions over the rotation, the translations, the variances and the latent, mean model. We implemented the entire model in the deep probabilistic programming language Pyro [7], using its automatic inference features. The results indicate that deep probabilistic programming readily allows the implementation, estimation and deployment of advanced non-Euclidean models relevant to structural bioinformatics. Specifically, we envision that THESEUS-PP can be used as a likelihood function in Bayesian protein structure prediction using deep probabilistic programming.

## II. METHODS

### A. Overall model

According to the THESEUS model [4], each observed protein structure **X**_*n*_ is a noisy observation of a rotated and translated latent, mean structure **M** with noise **E**_*n*_,

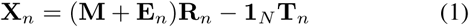

where *n* is an index that identifies the protein, **R** is a rotation matrix, **T** is a three-dimensional translation, **E** is the error and **M** and **X** are matrices with the atomic coordinate vectors along the rows,

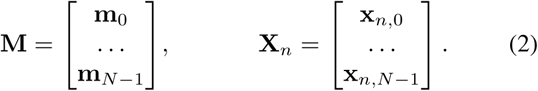

Another way of representing the model is seeing **X**_*n*_ as distributed according to a matrix-normal distribution with mean **M** and covariance matrices **U** and **V**-one concerning the rows and the other the columns.

The matrix-normal distribution can be considered as an extension of the standard multivariate normal distribution from vector-valued to matrix-valued random variables. Consider a random variable **X** distributed according to a matrix-normal distribution with mean **M**, which in our case is an *N*×3 matrix where N is the number of atoms. In this case, the matrix-normal distribution is further characterized by an *N* × *N* row covariance matrix **U** and a 3 × 3 column covariance **V**. Then, **X** ∼*ℳ 𝒩* (**M**, **U**, **V**) will be equal to

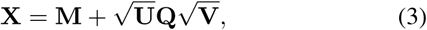

where **Q** is an *N* × 3 matrix with elements distributed according to the standard normal distribution.

To ensure identifiability, one (arbitrary) protein **X**_1_ is assumed to be a fixed noisy observation of the structure **M**:

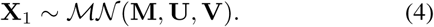

The other protein **X**_2_ is assumed to be a noisy observation of the rotated as well as translated mean structure **M**:

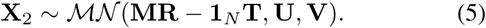

Thus, the model uses the same covariance matrices **U** and **V** for the matrix-normal distributions of both **X**_1_ and **X**_2_.

### B. Bayesian posterior

The graphical model of THESEUS-PP is shown in Figure 1. The corresponding Bayesian posterior distribution is

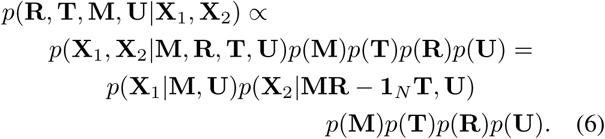

**Fig. 1:**
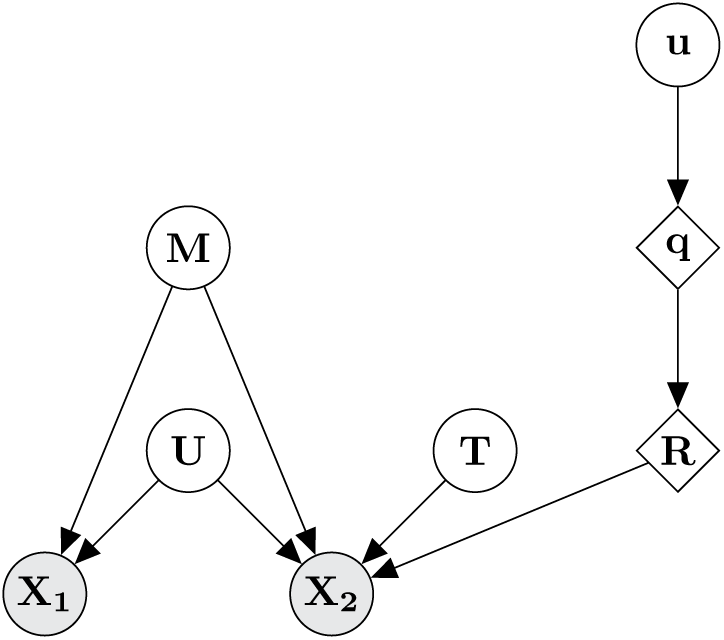
The THESEUS-PP model as a Bayesian graphical model. **M** is the latent, mean structure, which is an *N*-by-3 coordinate matrix, where *N* is the number of atoms. **T** is the translation. **q** is a unit quaternion calculated from three random variables **u** sampled from the unit interval. **R** is the corresponding rotation matrix. **U** is the among-row variance matrix of a matrix-normal distribution. **X**_1_ and **X**_2_ are *N*-by-3 coordinate matrices representing the proteins to be superimposed. Circles denote random variables. A lozenge denotes a deterministic transformation of a random variable. Shaded circles denote observed variables. Bold capital and bold small letters represent matrices and vectors, respectively.

Below, we specify how each of the priors and the likelihood function is formulated and implemented.

### C. Prior for the mean structure

Recall that according to the THESEUS-PP model, the atoms of the structures to be superimposed are noisy observations of a mean structure **M**. Typically, only *C*_*α*_ atoms are considered and in that case, *N* corresponds to the number of amino acids. Hence, we need to formulate a prior distribution over the latent, mean structure **M**.

We use an uninformative prior for **M**. Each element of **M** is sampled from a Student’s t-distribution with degrees of freedom (*v* = 1), mean (***µ*** = 0) and a uniform diagonal variance (***σ***^2^ = 3). The Student’s t-distribution is chosen over the normal distribution for reasons of numerical stability: the fatter tails of the Student’s t-distribution avoid numerical problems associated with highly variable regions.

### D. Prior over the rotation

In the general case, we have no *a priori* information on the optimal rotation. Hence, we use a uniform prior over the space of rotations. There are several ways to construct such a uniform prior. We have chosen a method that makes use of quaternions [8]. Quaternions are the 4-dimensional extensions of the better known 2-dimensional complex numbers. Unit quaternions form a convenient way to represent rotation matrices. For our goal, the overall idea is to sample uniformly from the space of unit quaternions. Subsequently, the sampled unit quaternions are transformed into the corresponding rotation matrices, which establishes a uniform prior over rotations.

A unit quaternion **q** = (*w, x, y, z*) is sampled in the following way. First, three independent random variables are sampled from the unit interval,

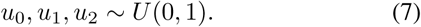

Then, four auxiliary deterministic variables (*θ*_1_, *θ*_2_, *r*_1_, *r*_2_) are calculated from *u*_1_, *u*_2_, *u*_3_,

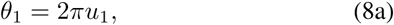

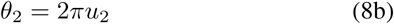

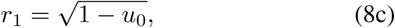

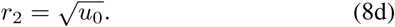

The unit quaternion **q** is then obtained in the following way,

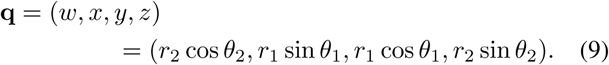

Finally, the unit quaternion **q** is transformed into its corresponding rotation matrix **R** as follows,

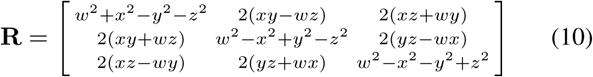

### E. Prior over the translation

For the translation, we use a standard trivariate normal distribution,

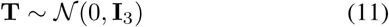

where **I**_3_ is the three-dimensional identity matrix.

### F. Prior over U

The Student’s t-distribution variance over the rows is sampled from the half-normal distribution with standard deviation set to 1.

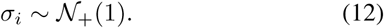

### G. Likelihood

In our case, the matrix-normal likelihood of THESEUS reduces to a product of univariate Student’s t-distributions. Again, we use the Student’s t-distribution rather than the normal distribution (as in THESEUS) for reasons of numerical stability. Below, we have used trivariate Student’s t-distributions with diagonal covariance matrices for ease of notation. The likelihood can thus be written as

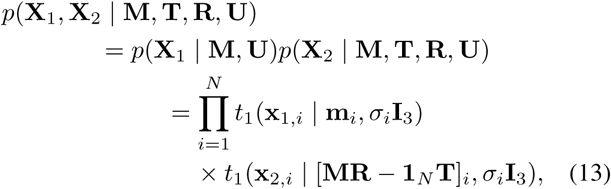

where the product runs over the matrix rows that contain the *x, y, z* coordinates of **X**_1_, **X**_2_ and the rotated and translated latent, mean structure **M**.

### H. Algorithm

#### Algorithm 1 The Theseus-PP model

►*Prior over the elements of* **M**

**m**_*i*_ ∼ *t*_1_(**0**, *σ*_*M*_ **I**_3_), where *i* indicates the atom position

►*Prior over the translation*

**T** ∼*𝒩* (**0**, **I**_3_)

►*Prior over rotation*

**u**_*j*_ ∼*U* [0, 1], for *j* from 0 to 2

**q** ← Quaternion(**u**)

**R** ← RotationMatrix(**q**)

► *Prior over diagonal covariance matrix* **U**

*σ*_*i*_ ∼*𝒩*_+_(1)

►*Likelihood over the N atom coordinates*

**x**_1,*i*_ ∼ *t*_1_(**m**_*i*_, *σ*_*i*_**I**_3_))

**x**_2,*i*_ ∼ *t*_1_([**RM −1**_*N*_ **T**]_*i*_, *σ*_*i*_**I**_3_)

### I. Initialization

Convergence of the MAP estimation can be greatly improved by selecting suitable starting values for certain variables and by transforming the two structures **X**_1_ and **X**_2_ in a suitable way. First, we pre-superimpose the two structures using conventional least-squares superposition. Therefore, the starting rotation can be initialized close to the identity matrix (ie., no rotation). This is done by setting the vector **u** to (0.9, 0.1, 0.9).

We further improve performance by initializing the mean structure **M** to the average of the two pre-superimposed structures **X**_1_ and **X**_2_.

### J. Maximum a-posteriori optimization

We performed MAP estimation using Pyro’s AutoDelta guide. For optimization, we used AdagradRMSProp [9], [10] with the default parameters for the learning rate (1.0), momentum (0.1) and step size modulator (1.0×10^-16^). A fragment of the model implementation in Pyro can be seen in Figure 3 in the Appendix.

Convergence was detected using Earlystop from Pytorch’s Ignite library (version 0.2.0) [11]. This method evaluates the stabilization of the error loss and stops the optimization according to the value of the *patience* parameter. The *patience* value was set to 25.

## III. MATERIALS

### Proteins

The algorithm was tested on several proteins from the RCSB protein database [12] that were obtained from Nuclear Magnetic Resonance (NMR) experiments. Such structures typically contain several models of the same protein. These models represent the structural dynamics of the protein in an aqueous medium and thus typically contain both conserved and variable regions. This makes them challenging targets for conventional RMSD superposition. We used the following structures: 1ADZ, 1AHL, 1AK7, 2CPD, 2KHI, 2LKL and 2YS9.

## IV. RESULTS

The algorithm was executed 15 times on each protein (see TABLE I) with different seeds. The computations where carried on a Intel Core i7-8750H CPU 2.20GHz processor.

**TABLE I:** Results of applying THESEUS-PP to the test structures. First column: PDB identifier. Second column: the number of *C*_*α*_ atoms used in the superposition. Third column: the model identifiers. Fourth column: mean convergence time and standard deviation. Last column: Number of epochs.

| PBD ID | Length<br>(Amino Acids) | Protein<br>Models | Average<br>Computational Time<br>(seconds) | Epochs |
| --- | --- | --- | --- | --- |
| 1ADZ | 71 | 0 and 1 | $0.64 \pm 0.15$ | $210 \pm 45$ |
| 1AHL | 49 | 0 and 2 | $0.53 \pm 0.119$ | $166 \pm 36$ |
| 1AK7 | 174 | 0 and 1 | $0.74 \pm 0.23$ | $222 \pm 67$ |
| 2CPD | 99 | 0 and 2 | $0.51 \pm 0.093$ | $161 \pm 30$ |
| 2KHI | 95 | 0 and 1 | $0.47 \pm 0.11$ | $149 \pm 39$ |
| 2LKL | 81 | 0 and 8 | $0.59 \pm 0.092$ | $187 \pm 30$ |
| 2YS9 | 70 | 0 and 3 | $0.47 \pm 0.11$ | $144 \pm 33$ |

An example of a pair of superimposed structures is shown in Figure 2. For comparison, the superposition resulting from the conventional RMSD method, as calculated using Biopython [13], is shown on the left (Figure 2a). The THESEUS-PP superposition is shown on the right (Figure 2b). Note how the former fails to adequately distinguish regions with high from regions with low variance, resulting in poor matching of conserved regions. Additional, similar figures of superimposed structures can be found in the Appendix.

**Fig. 2:**
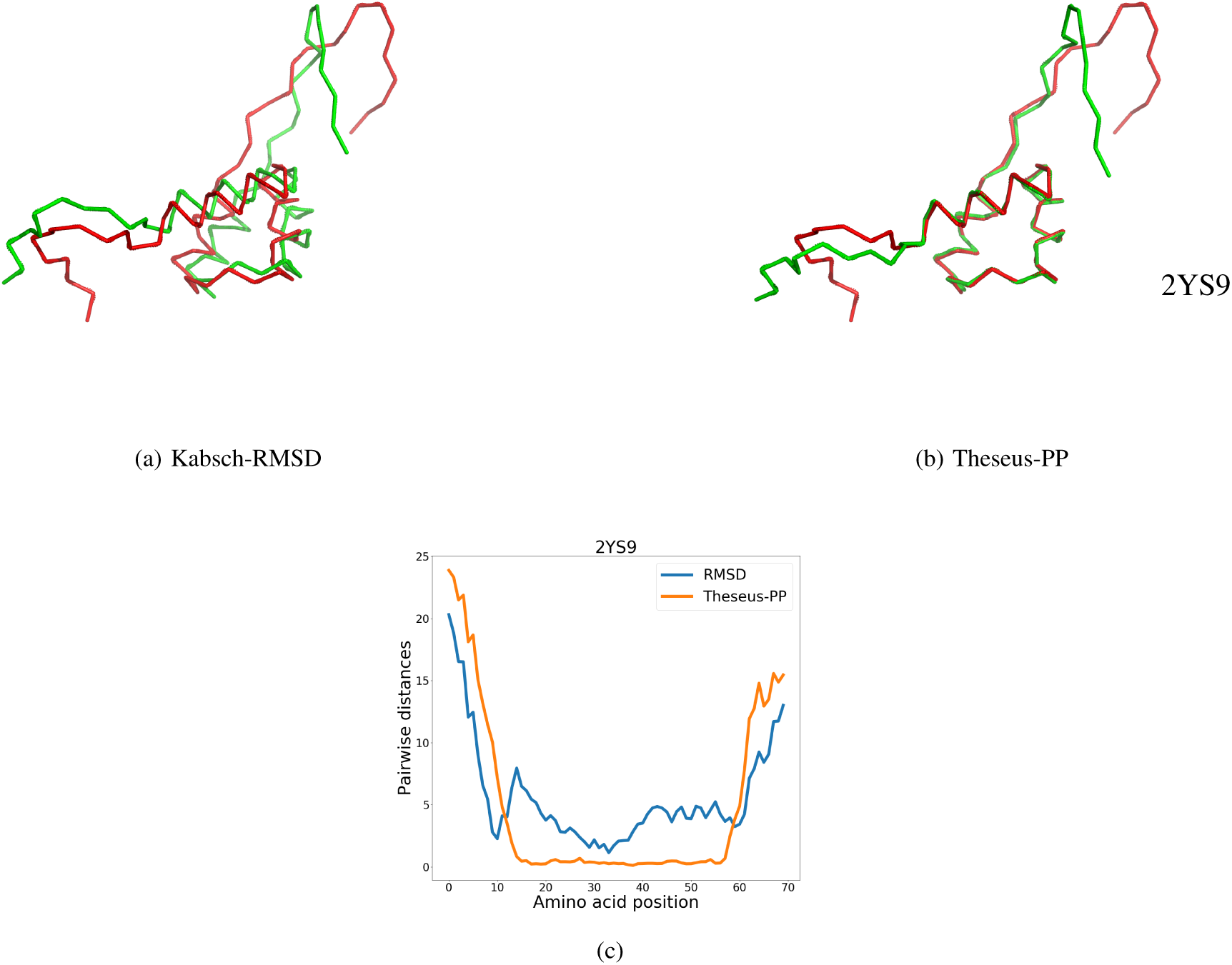
Protein superposition for two conformations of protein 2YS9 obtained from (a) conventional RMSD superimposition and (b) THESEUS-PP. The protein in green is rotated (*X*_2_). The images are generated with PyMOL [14]. Graph (c) shows the pairwise distances (in Å) between the *C*_*α*_ coordinates of the structure pairs. The blue and orange lines represent RMSD and THESEUS-PP superposition, respectively.

## V. CONCLUSION

Probabilistic programming is a powerful, emerging paradigm for probabilistic protein structure analysis, prediction and design. Here, we present a Bayesian model for protein structure superposition implemented in the deep probabilistic programming language Pyro and building on the previously reported THESEUS maximum likelihood model. MAP estimates of its parameters are readily obtained using Pyro’s automated inference engine.

The original THESEUS algorithm, which makes use of maximum likelihood estimation using iterative expectation maximization, is considerably faster with an average execution time under 0.1 s. Although some of the longer execution time in THESEUS-PP is due to the use of variational inference and priors, it is clear that the flexibility and productivity of a probabilistic programming language can come with a speed penalty.

Recently, end-to-end protein protein structure prediction using deep learning methods has become possible [15]. We envision that Bayesian protein structure prediction will soon be possible using a deep probabilistic programming approach, which will lead to protein structure predictions with associated statistical uncertainties. In order to achieve this goal, suitable error models and likelihood functions need to be developed and incorporated in these models. The THESEUS-PP model can potentially serve as such an error model, by interpreting **M** as the predicted structure and a single rotated and translated **X** as the observed protein structure. During training of the probabilistic model, regions in **M** that are wrongly predicted can be assigned high variance, while correctly predicted regions can be assigned low variance. Thus, it can be expected that an error model based on THESEUS-PP will make estimation of these models easier, as the error function can more readily distinguish between partly correct and entirely wrong predictions, which is notoriously difficult for RMSD-based methods [16].

## CONTRIBUTIONS AND ACKNOWLEDGEMENTS

Implemented algorithm in Pyro: *LSM*. Contributed code: *ASA, AM*. Wrote article: *LSM, TH, ASA*. Prototyped algorithm in the probabilistic programming language PyMC3 [17]: *TH, WB, BNR*. Performed experiments: *LSM*. Designed experiments: *TH, DT*. *LSM* and *ASA* acknowledge support from the Independent Research Fund Denmark (grant: “Resurrecting ancestral proteins in silico to understand how cancer drugs work”) and Innovationsfonden/Skanned.com (grant: “Intelligent accounting document management using probabilistic programming”), respectively. We thank Jotun Hein, Michael Golden, Kanti Mardia and Wouter Boomsma for suggestions and discussions.

## DATA AND SOFTWARE AVAILABILITY

Pyro [7] and PyTorch [11] based code is a available at https://github.com/LysSanzMoreta/Theseus-PP

## VI. APPENDIX

**Fig. 3:**
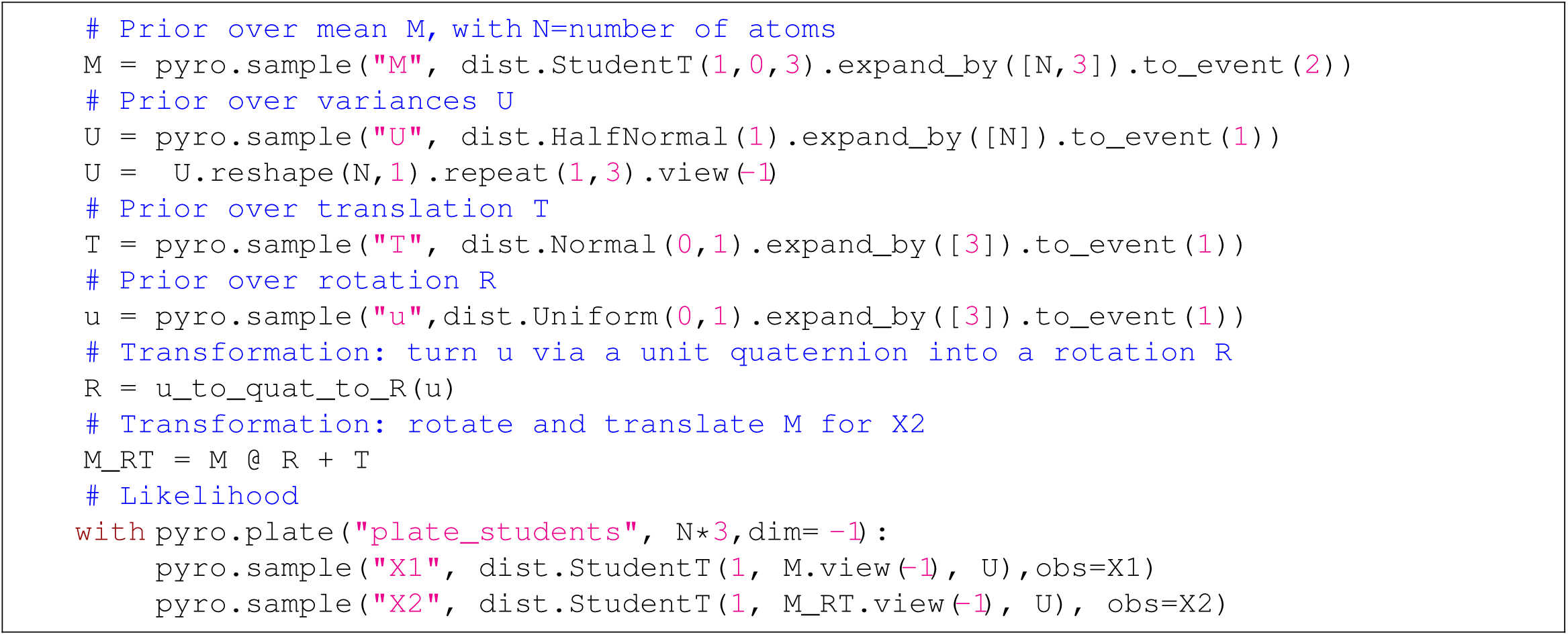
Code fragment from the THESEUS-PP implementation in Pyro. *pyro.sample* calls a primitive stochastic function from which a named sample is drawn. *expand_by* specifies the shape of the batch that is to be drawn from the distribution. *pyro.plate* declares the variables within a tensor dimension as conditionally independent, while *to_event* declares them as dependent.

**Fig. 4:**
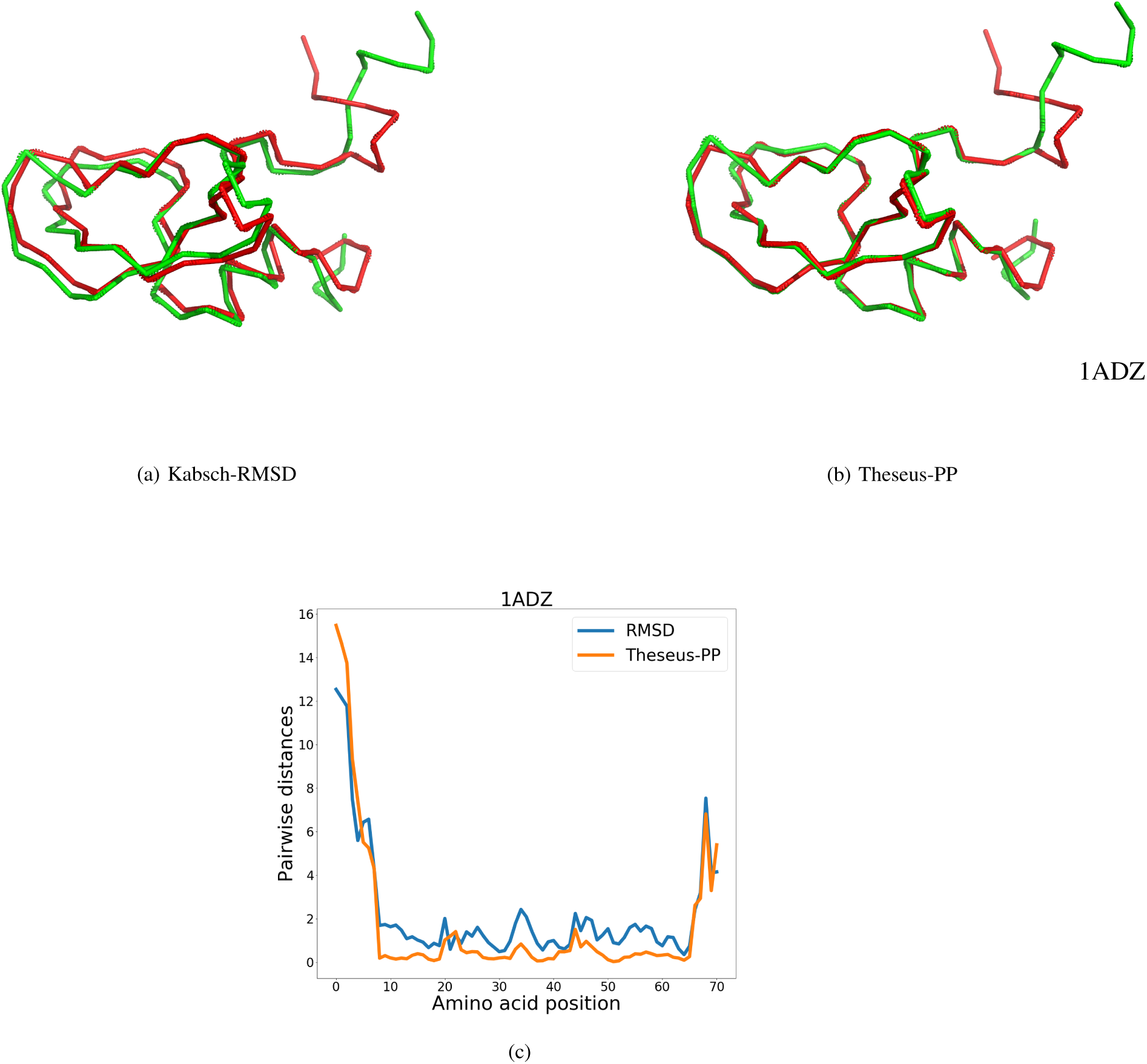
Protein superposition for two conformations of protein 1ADZ obtained from (a) conventional RMSD superimposition and (b) THESEUS-PP. The protein in green is rotated (*X*_2_). The images are generated with PyMOL [14]. Graph (c) shows the pairwise distances (in Å) between the *C*_*α*_ coordinates of the structure pairs. The blue and orange lines represent RMSD and THESEUS-PP superposition, respectively.

**Fig. 5:**
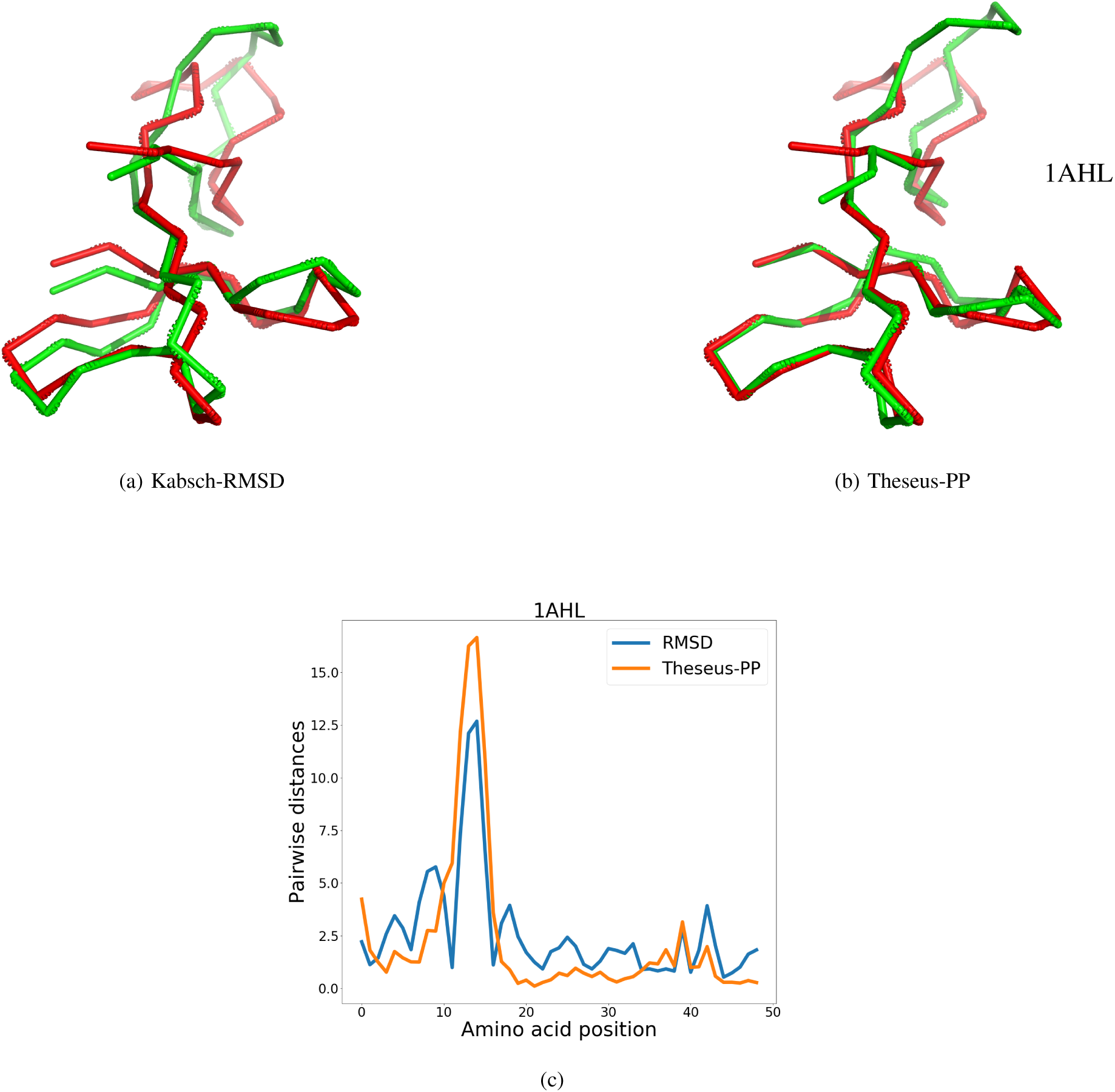
Protein superposition for two conformations of protein 1AHL obtained from (a) conventional RMSD superimposition and (b) THESEUS-PP. The protein in green is rotated (*X*_2_). The images are generated with PyMOL [14]. Graph (c) shows the pairwise distances (in Å) between the *C*_*α*_ coordinates of the structure pairs. The blue and orange lines represent RMSD and THESEUS-PP superposition, respectively.

**Fig. 6:**
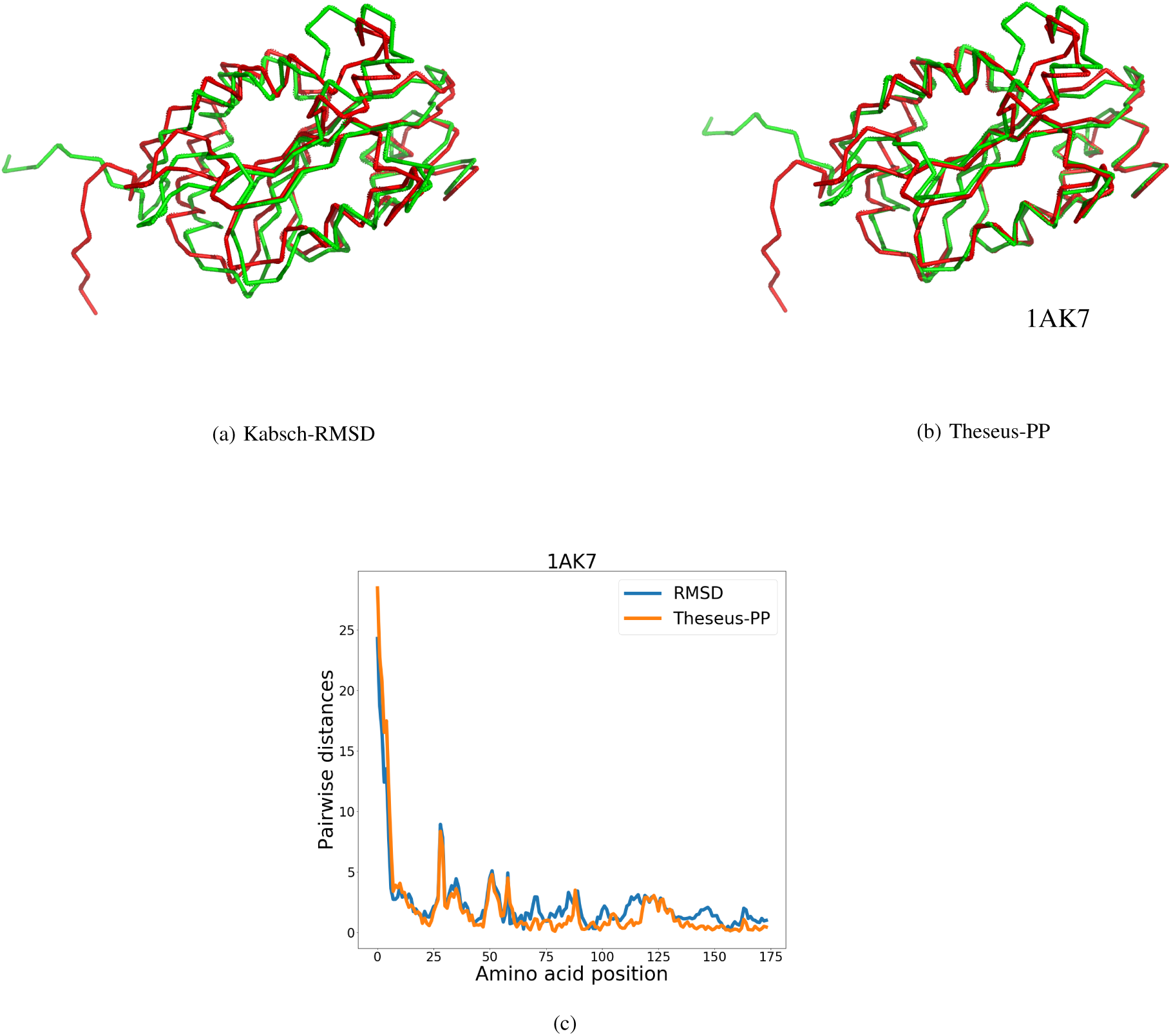
Protein superposition for two conformations of protein 1AK7 obtained from (a) conventional RMSD superimposition and (b) THESEUS-PP. The protein in green is rotated (*X*_2_). The images are generated with PyMOL [14]. Graph (c) shows the pairwise distances (in Å) between the *C*_*α*_ coordinates of the structure pairs. The blue and orange lines represent RMSD and THESEUS-PP superposition, respectively.

**Fig. 7:**
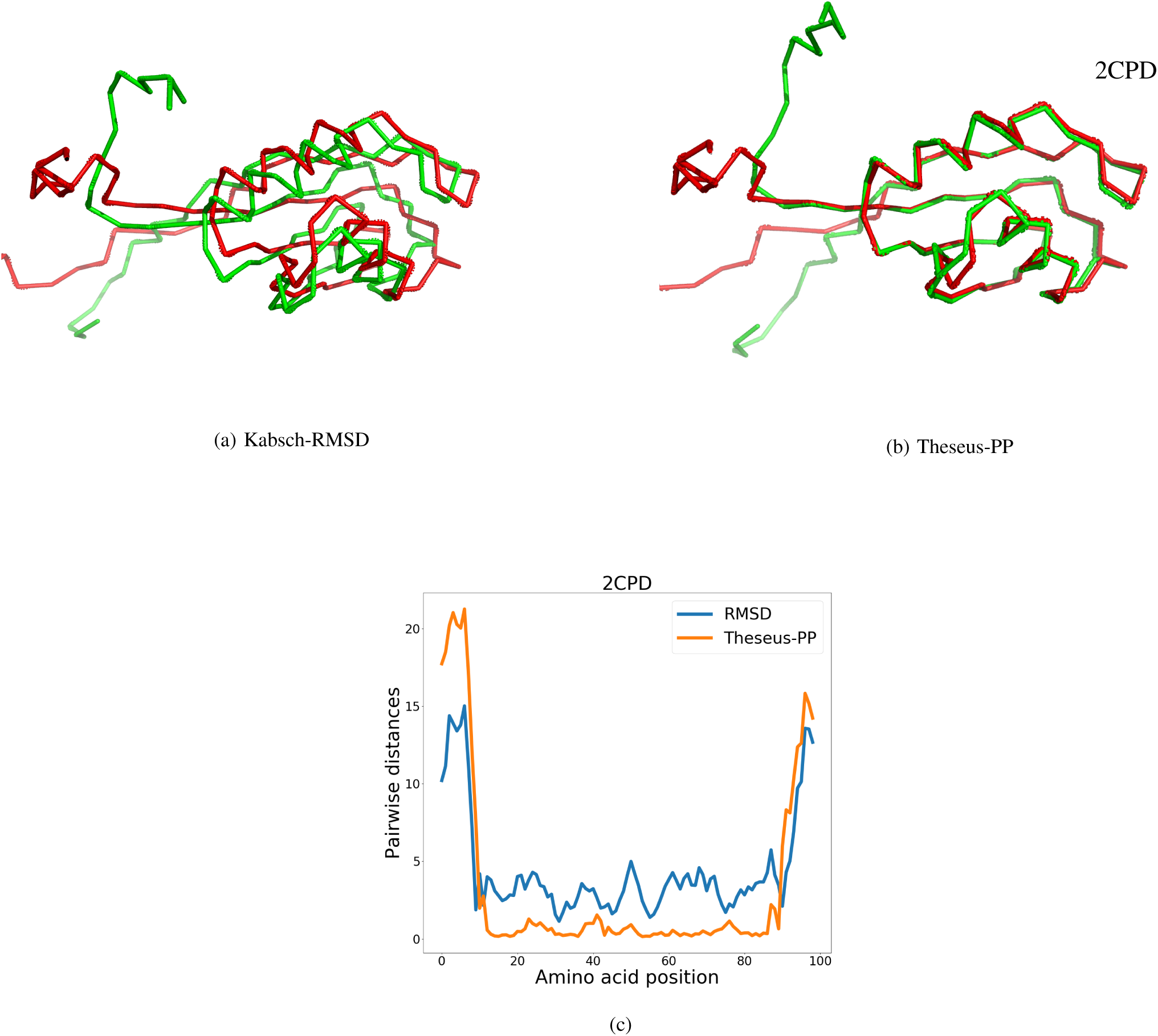
Protein superposition for two conformations of protein 2CPD obtained from (a) conventional RMSD superimposition and (b) THESEUS-PP. The protein in green is rotated (*X*_2_). The images are generated with PyMOL [14]. Graph (c) shows the pairwise distances (in Å) between the *C*_*α*_ coordinates of the structure pairs. The blue and orange lines represent RMSD and THESEUS-PP superposition, respectively.

**Fig. 8:**
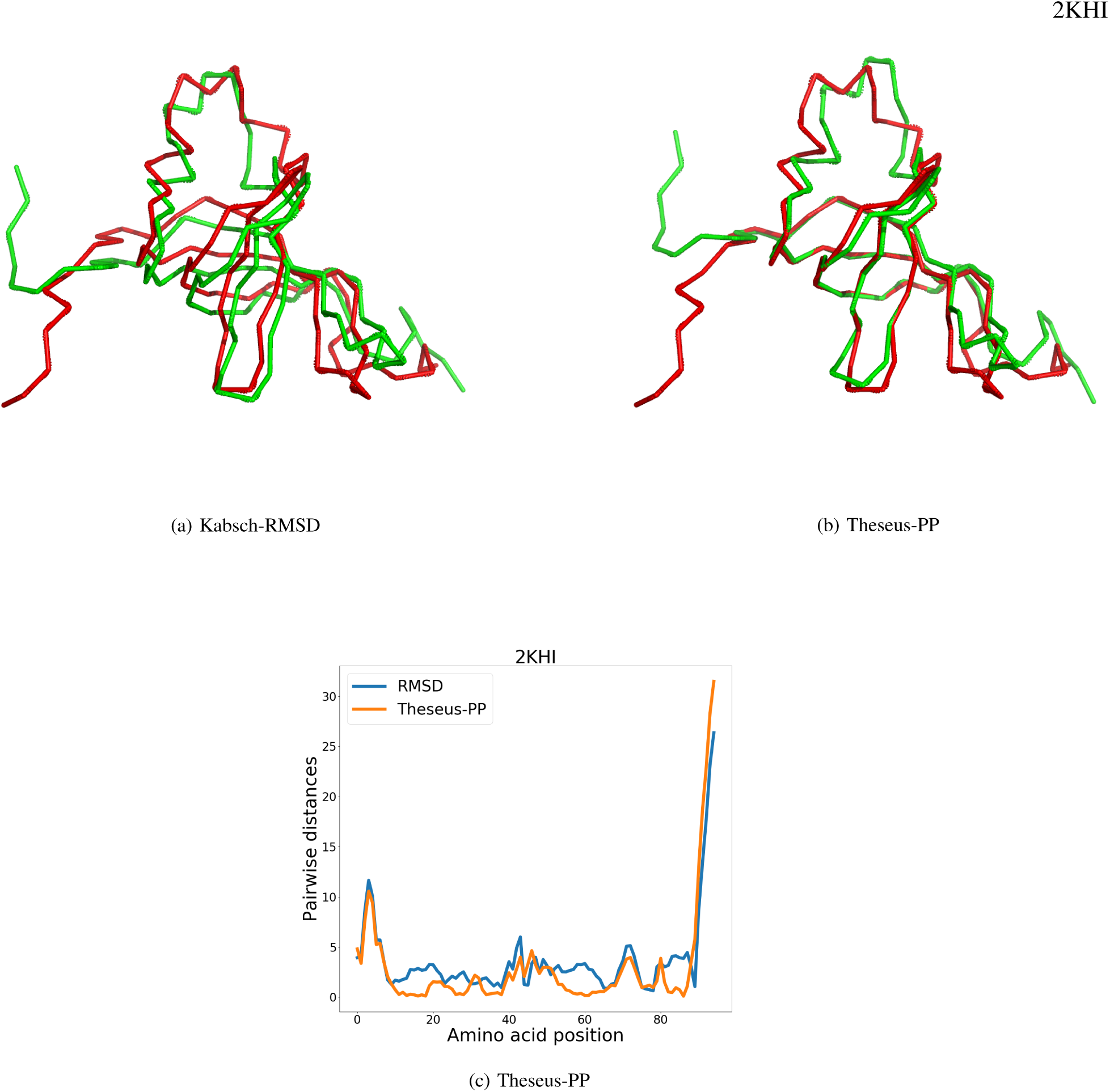
Protein superposition for two conformations of protein 2KHI obtained from (a) conventional RMSD superimposition and (b) THESEUS-PP. The protein in green is rotated (*X*_2_). The images are generated with PyMOL [14]. Graph (c) shows the pairwise distances (in Å) between the *C*_*α*_ coordinates of the structure pairs. The blue and orange lines represent RMSD and THESEUS-PP superposition, respectively.

**Fig. 9:**
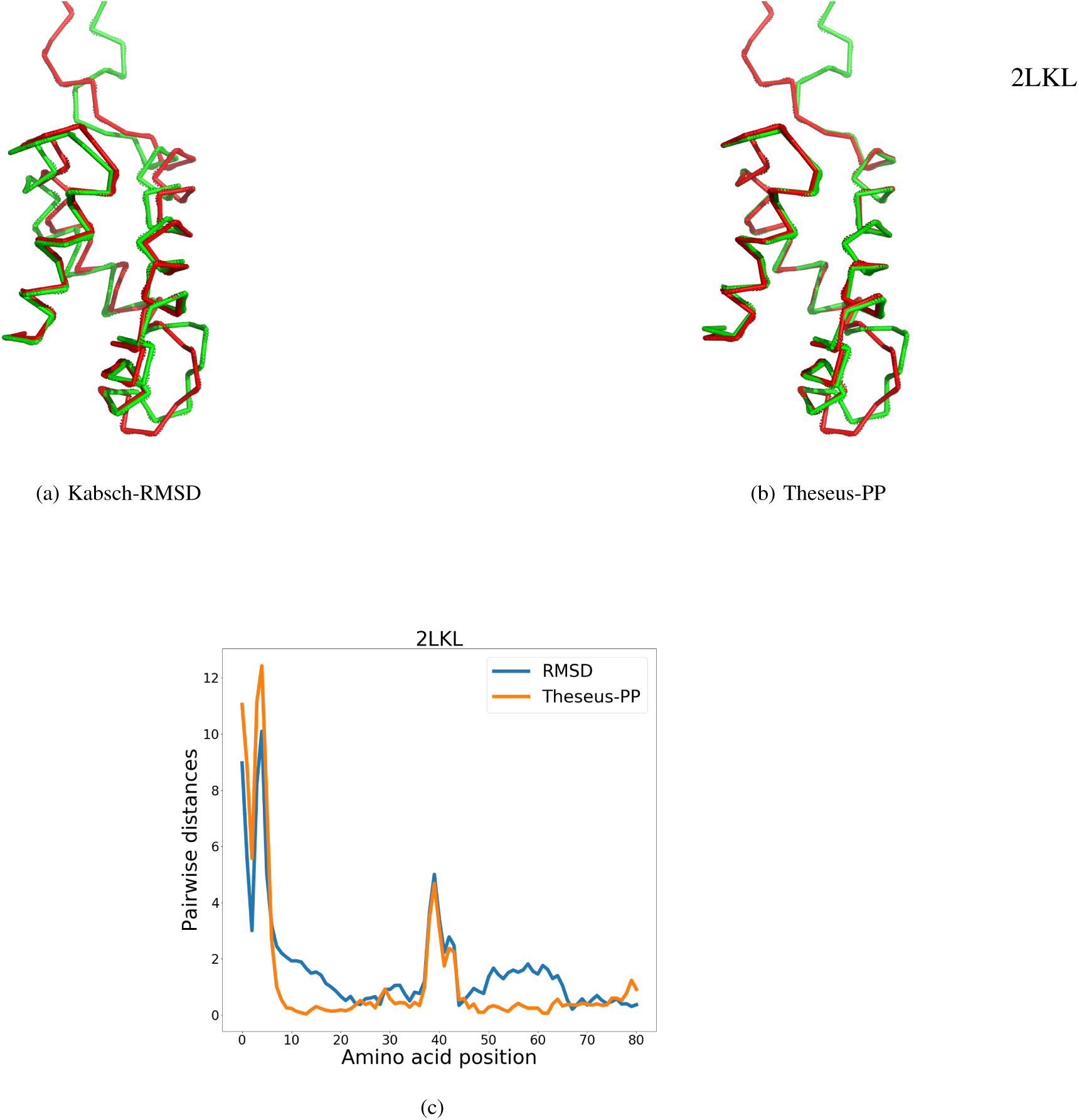
Protein superposition for two conformations of protein 2LKL obtained from (a) conventional RMSD superimposition and (b) THESEUS-PP. The protein in green is rotated (*X*_2_). The images are generated with PyMOL [14]. Graph (c) shows the pairwise distances (in Å) between the *C*_*α*_ coordinates of the structure pairs. The blue and orange lines represent RMSD and THESEUS-PP superposition, respectively.

## Notes

#### Summary of Updates

The model previously presented in this paper has been updated and improved notoriously. Mainly the model has been simplified by removing one of the translations and stabilized by changing some of the priors distributions. These changes have subsequently ameliorated the speed and the accuracy of the model.

